# Zoonotic spillover of SARS-CoV-2: mink-adapted virus in humans

**DOI:** 10.1101/2021.03.05.433713

**Authors:** Lukasz Rabalski, Maciej Kosinski, Natalia Mazur-Panasiuk, Boguslaw Szewczyk, Krystyna Bienkowska-Szewczyk, Ravi Kant, Tarja Sironen, Krysztof Pyrć, Maciej Grzybek

**Author notes:** Corresponding authors: Maciej Grzybek and Krzysztof Pyrc Corresponding author emails: ma &.

## Abstract

The COVID-19 pandemic caused by SARS-CoV-2 started in fall 2019. A range of different mammalian species, including farmed mink, have been confirmed as susceptible to infection with this virus. We report here the spillover of mink-adapted SARS-CoV-2 from farmed mink to humans after extensive adaptation that lasted at least 3 months. We found the presence of four mutations in the S gene (that gave rise to variant: G75V, M177T, Y453F and C1247F) and others in an isolate obtained from SARS-CoV-2 positive patient.

## 1. Introduction

Coronaviruses are known as potential zoonotic pathogens, and severe acute respiratory syndrome coronavirus 2 (SARS-CoV-2) is the third highly pathogenic member of this family to have emerged in the 21^st^ century (1). While mass vaccinations are currently underway, the question of the fate of the virus remains open. The concept of herd immunity and eradication of the virus is somewhat unrealistic when considering the prevalence, genetic diversity, and the existing animal reservoirs. Thus far, SARS-CoV-2 infections have been reported in different mammalian species worldwide, including dogs, cats, tigers, lions, ferrets, minks, felines, and deer (2,3). SARS-CoV-2 infections in farmed mink have recently been confirmed in European countries (4). Transmission of the virus from infected mink to humans has been reported in Denmark and the Netherlands (5–7). After Denmark, Poland is the second-largest producer of mink pelts in Europe, with 354 active Polish mink farms harbouring approximately 6.3 million mink in total (8).

## 2. Material and methods

This study was approved by the Independent Bioethical Committee for Scientific Research at the Medical University of Gdansk, Gdansk, Poland (Statement no. NKBBN/183/2020).

SARS-CoV-2 genome sequencing was performed at the University of Gdansk, Poland using sample containing RNA isolated from positive swab (amplification of two target genes in RT-PCR). ARTICv3 amplicon generation followed by Oxford Nanopore Technology MinION run was performed (Quick 2020). Reads were basecalled, debarcoded and trimmed to delete adapter, barcode and PCR primer sequences. ARCTIC pipeline software were used to generate SARS-CoV-2 genome. Phylogenetic analysis was performed using the procedure recommended by Nextstrain.org (Hadfield et al., 2018)

## 3. Results and Discussion

The capacity of coronaviruses to transfer between mammalian species is not surprising. First, the transfer of highly pathogenic variants from bats usually occurs via intermediate hosts. Infection of dromedary camels and palm civets and later transfer to humans has been described for MERS-CoV and SARS-CoV, respectively (9). A more comprehensive analysis of existing strains suggests that this was not an exception. The human coronavirus OC43 virus (HCoV-OC43) is a betacoronavirus described for the first time in the 1960s. This pathogen is associated with upper and lower respiratory tract disease in humans (10). Interestingly, closely related species are found in cattle (bovine coronavirus) and dogs (canine respiratory coronavirus). While the exact transfer route between different species remains unknown, it is conceivable that as in SARS-CoV-2, the virus has jumped between humans, and companion and farmed animals in the past (11).

Recently, we and others have detected SARS-CoV-2 infection in farmed minks in Northern Poland (12,13). The first report identified SARS-CoV-2 in samples collected from mink in mid-November 2020. While the prevalence of the virus was low and, most likely, only isolated cases were present, we have sequenced the isolates and have shown that the positive signal did not originate from contamination. The most recent data collected and deposited by the laboratory of the National Veterinary Research Institute in Poland has indicated isolation of the SARS-CoV-2 variants from animals at the same farm. Phylogenetic analysis of the data have indicated that the virus belongs to the B.1.1.279 lineage (Pangolin classification), which is not surprising considering the prevalence of this variant in Europe. Genome analysis shows that the new isolates carry the combination of mutations typical of viruses isolated already in November 2020 from mink (listed in Figure 1 bottom left radial tree), but additional new changes have accumulated since then. These include the Y453F mutation, which was previously reported to have emerged in minks during serial passages (e.g., in Denmark and recently Lithuania), and a novel mutation not present in any global SARS-CoV-2 isolate that truncate ORF 7b at position L22. With all the available data taken into account, we speculate that the virus was present already in the mink population by November 2020, presumably after a single introduction during the late summer or autumn of that year.

**Figure 1.**
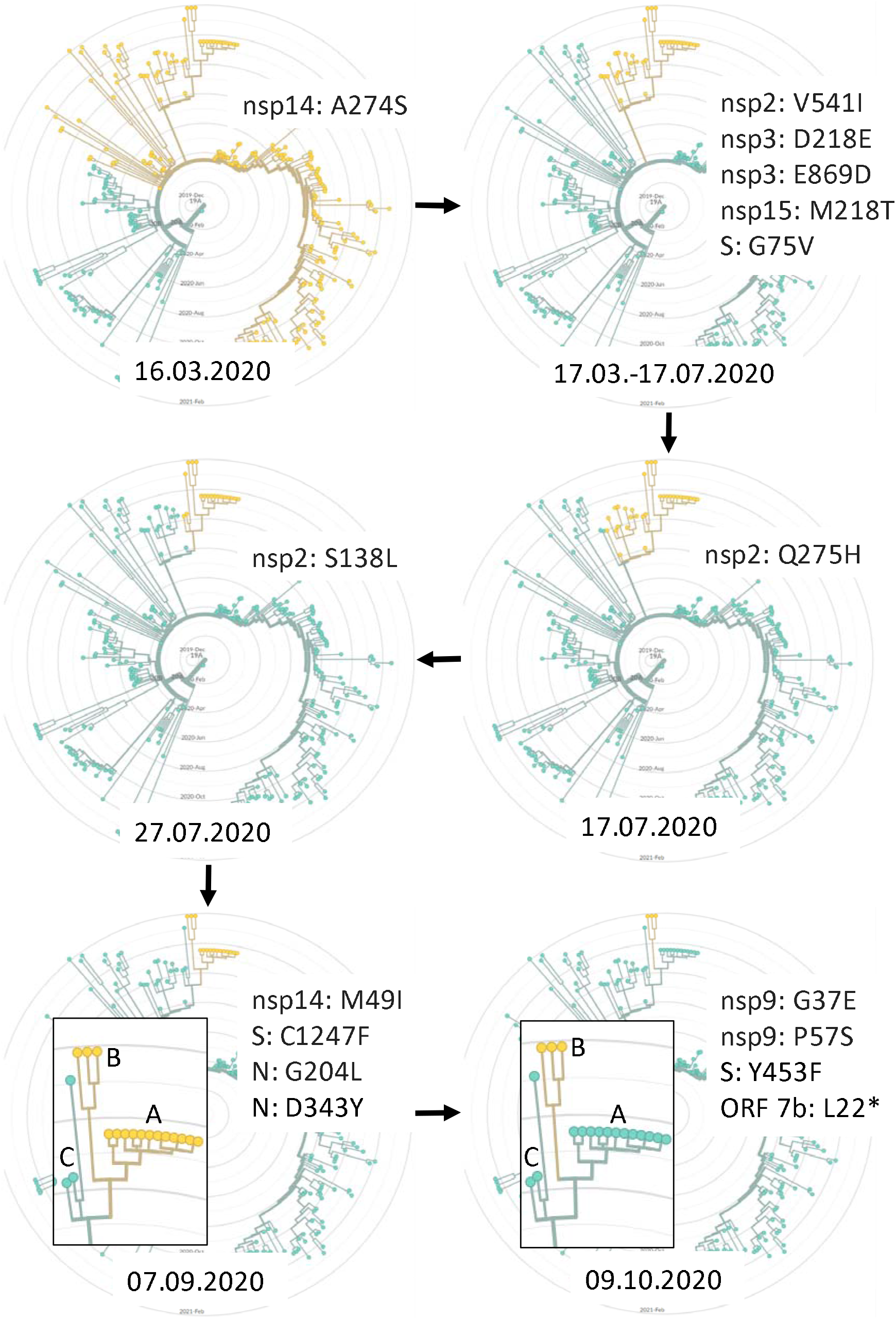
Phylogenetic analysis of SARS-CoV-2 lineage B.1.1.279 combined with inferred time (bottom of each radial time tree) of fixing mutations (upper right of each radial time tree) in all isolates form a group that leads to the generation of the mink variants. Yellow colour represents new variants, A - November 2020 mink isolates, B - January 2021 mink isolates and single human isolate, C - nearest neighbours human isolates that share a common ancestor with A-B: Norway/4235/2020, Germany/NW-HHU-340/2020, Iceland/4563/2021.

In the current study, we have identified a case of infection with a mink-adapted variant in humans. Following the identification of SARS-CoV-2 cases in farmed animals, the exposed staff were tested for SARS-CoV-2 (nasopharyngeal swabs) using RT-qPCR. A single positive case was detected in a sample collected on 1^st^ February 2021 and was sequenced using the ARTIC Nanopore technology protocol. The resulting sequence has been deposited in GISAID under the accession number EPI_ISL_1034274. The host was asymptomatic. Phylogenetic analysis (Figure 1) shows that the virus clusters closely with viruses isolated from mink (group B on bottom radial trees). Further, mutations characteristic of the mink-adapted variant were present in the virus's genomic sequence (Table 1), confirming that it was most likely contracted by the infected subject from animals.

**Table 1.** Representation of mutations in mink originated SARS-CoV-2 isolates. Group A - November 2020 mink isolates, Group B - January 2021 mink isolates and single human isolate, C - nearest neighbours in lineage B.1.1.279 human isolates. The yellow colour represents mutations fixed in November 2020. The red colour represents mutations acquired during three months of passage at the mink farm.

| nt position | aa variant | Reference | Group A | Group B | Group C |
| --- | --- | --- | --- | --- | --- |
| 1218 | nsp2: S138L | C | T | T | T |
| 1630 | nsp2: Q275H | A | T | T | T |
| 2426 | nsp2: V541I | G | A | A | A |
| 3373 | nsp3: D218E | C | A | A | A |
| 5326 | nsp3: E869D | G | T | T | T |
| 12795 | nsp9: G37E | G | G | A | G |
| 12854 | nsp9: P57S | C | C | T | C |
| 14408 | nsp12: P323L | C | T | T | T |
| 18186 | nsp14: M49I | G | T | T | G |
| 18859 | nsp14: A274S | G | T | T | T |
| 20273 | nsp15: M218T | T | C | C | C |
| 21786 | S: G75V | G | T | T | T |
| 22092 | S: M177T | T | T | C | T |
| 22920 | S: Y453F | A | A | T | A |
| 25302 | S: C1247F | G | T | T | G |
| 27820 | ORF 7b: L22* | T | T | A | T |
| 28881/2 | N: R203K | GG | AA | AA | AA |
| 28884 | N: G204L | G | T | T | G |
| 29300 | N: D343Y | G | T | T | G |

The emerging variant clearly reflects adaptation of the virus to the animal host, with some point mutations fixed early during transmission between animals and additional changes accumulating over time (Figure 1, bottom right). While the mutations' exact role is to be determined, they may provide increased fitness in the new host (14). It remains unknown whether these features alter the course of disease transmissibility, or immunogenicity in humans (15). However, the variant should be tracked in the general population. Our results confirm the need for country-scale epizootiological monitoring and careful analysis of SARS-CoV-2 positive patients.

## Data availability

The complete genome sequences of SARS-CoV-2 isolated from the patient was deposited in GISAID under the accession number: EPI_ISL_1034274

## Authors’ contributions

The study was conceived and designed by LR and MG. Data handling: LR, MK, MG, KP. Material handling and laboratory work: LR, MK, NMP. Statistical and phylogenetic analysis was carried by LR. Interpretation of data: LR, KP, MK. Data visualisation: LR. Supervision: MG, KP, TS. The manuscript was written by LR, MG, KP, RK, TS, BS, KBS in consultation with all co-authors. MG, KP, LR, TS, RK revised the manuscript. All authors accepted the final manuscript version.

## Acknowledgements

We gratefully acknowledge the Authors from the Originating laboratories responsible for obtaining the specimens and the Submitting laboratories where genetic sequence data were generated and shared via the GISAID Initiative, on which this research is partially based.

## Notes

**Conflict of interest**: None declared

### Competing Interest Statement

The authors have declared no competing interest.

## REFERENCES

1. Cucinotta D, Vanelli M. WHO Declares COVID-19 a Pandemic. Acta Biomed [Internet]. 2020;91(1):157–60. Available from: http://www.ncbi.nlm.nih.gov/pubmed/32191675

2. Palmer M V, Martins M, Falkenberg S, Buckley A, Caserta LC, Mitchell PK, et al. Susceptibility of white-tailed deer (Odocoileus virginianus) to SARS-CoV-2. bioRxiv [Internet]. 2021 Jan 1;2021.01.13.426628. Available from: http://biorxiv.org/content/early/2021/01/14/2021.01.13.426628.abstract

3. Abdel-Moneim AS, Abdelwhab EM. Evidence for SARS-CoV-2 Infection of Animal Hosts. Pathogens [Internet]. 2020 Jun 30;9(7):529. Available from: https://www.mdpi.com/2076-0817/9/7/529

4. World Organisation for Animal Health. World Organisation for Animal Health [Internet]. 2021 [cited 2021 Jan 12]. Available from: https://www.oie.int/en/scientific-expertise/specific-information-and-recommendations/questions-and-answers-on-2019novel-coronavirus/events-in-animals/

5. Hammer AS, Quaade ML, Rasmussen TB, Fonager J, Rasmussen M, Mundbjerg K, et al. SARS-CoV-2 Transmission between Mink (Neovison vison) and Humans, Denmark. Emerg Infect Dis [Internet]. 2021 Feb;27(2). Available from: http://www.nc.cdc.gov/eid/article/27/2/20-3794_article.htm

6. Koopmans M. SARS-CoV-2 and the human-animal interface: outbreaks on mink farms. Lancet Infect Dis [Internet]. 2020 Nov;1. Available from: https://linkinghub.elsevier.com/retrieve/pii/S1473309920309129

7. Oude Munnink BB, Sikkema RS, Nieuwenhuijse DF, Molenaar RJ, Munger E, Molenkamp R, et al. Transmission of SARS-CoV-2 on mink farms between humans and mink and back to humans. Science (80-) [Internet]. 2020 Nov 10;eabe5901. Available from: https://www.sciencemag.org/lookup/doi/10.1126/science.abe5901

8. EU Fur Association. EU Fur Association [Internet]. 2020 [cited 2020 Dec 13]. Available from: https://www.sustainablefur.com/

9. de Wit E, van Doremalen N, Falzarano D, Munster VJ. SARS and MERS: recent insights into emerging coronaviruses. Nat Rev Microbiol [Internet]. 2016 Aug 27;14(8):523–34. Available from: http://www.nature.com/articles/nrmicro.2016.81

10. Kahn JS, McIntosh K. History and Recent Advances in Coronavirus Discovery. Pediatr Infect Dis J [Internet]. 2005 Nov;24(11):S223–7. Available from: https://journals.lww.com/00006454-200511001-00012

11. Szczepanski A, Owczarek K, Bzowska M, Gula K, Drebot I, Ochman M, et al. Canine Respiratory Coronavirus, Bovine Coronavirus, and Human Coronavirus OC43: Receptors and Attachment Factors. Viruses [Internet]. 2019 Apr 5;11(4):328. Available from: https://www.mdpi.com/1999-4915/11/4/328

12. Rabalski L, Kosinski M, Smura T, Aaltonen K, Kant R, Sironen T, et al. Detection and molecular characterisation of SARS-CoV-2 in farmed mink (Neovison vison) in Poland. bioRxiv. 2020;

13. GISAID. GISAID [Internet]. 2021 [cited 2021 Feb 18]. Available from: https://www.epicov.org/epi3/EPI_ISL_984305,EPI_ISL_984307

14. Zhao J, Cui W, Tian B. The Potential Intermediate Hosts for SARS-CoV-2. Front Microbiol [Internet]. 2020 Sep 30;11. Available from: https://www.frontiersin.org/article/10.3389/fmicb.2020.580137/full

15. Hayashi T, Yaegashi N, Konishi I. Effect of RBD mutation (Y453F) in spike glycoprotein of SARS-CoV-2 on neutralizing antibody affinity. bioRxiv [Internet]. 2020 Jan 1;2020.11.27.401893. Available from: http://biorxiv.org/content/early/2020/11/28/2020.11.27.401893.abstract

